# Multi-compartment analysis of the complex gradient-echo signal quantifies myelin breakdown in premanifest Huntington’s disease

**DOI:** 10.1101/2021.03.12.435119

**Authors:** Chiara Casella, Elena Kleban, Anne E. Rosser, Elizabeth Coulthard, Hugh Rickards, Fabrizio Fasano, Claudia Metzler-Baddeley, Derek K. Jones

## Abstract

White matter (WM) alterations have been identified as a relevant pathological feature of Huntington’s disease (HD). Increasing evidence suggests that WM changes in this disorder are due to alterations in myelin-associated biological processes. Multi-compartmental analysis of the complex gradient-echo MRI signal evolution in WM has been shown to quantify myelin *in vivo*, therefore pointing to the potential of this technique for the study of WM myelin changes in health and disease. This study first characterized the reproducibility of metrics derived from the complex multi-echo gradient-recalled echo (mGRE) signal across the corpus callosum in healthy participants, finding highest reproducibility in the posterior callosal segment. Subsequently, the same analysis pipeline was applied in this callosal region in a sample of premanifest HD patients (n = 19) and age, sex and education matched healthy controls (n = 21). In particular, we focused on two myelin-associated derivatives: i. the myelin water signal fraction (f_m_), a parameter dependent on myelin content; and ii. the difference in frequency between myelin and intra-axonal water pools (Δω), a parameter dependent on the ratio between the inner and the outer axonal radii. f_m_ was found to be lower in HD patients (β = −0.13, p = 0.03), while Δω did not show a group effect. Performance in tests of working memory, executive function, social cognition and movement was also assessed, and a greater age-related decline in executive function was detected in HD patients (β = −0.06, p = 0.006), replicating previous evidence of executive dysfunction in HD. Finally, the correlation between f_m_, executive function, and proximity to disease onset was explored in patients, and a positive correlation between executive function and f_m_ was detected (r = 0.542; p = 0.02). This study emphasises the potential of complex mGRE signal analysis for aiding understanding of HD pathogenesis and progression. Moreover, expanding on evidence from pathology and animal studies, it provides novel *in vivo* evidence supporting myelin breakdown as an early feature of HD.

## 1. Introduction

### 1.1. Why study myelin changes in Huntington’s disease?

Huntington’s disease (HD) is a debilitating genetic disorder caused by an expansion of the CAG (cytosine, adenine, guanine) repeat within the huntingtin (*HTT*) gene, and characterised by motor, cognitive and psychiatric symptoms associated with neuropathological decline. Although clinical onset of the disease is formally identified with the development of movement symptoms (Nopoulos, 2016), critical pathogenic events are present early on in the disease course (see Casella et al., 2020 for a review). Accordingly, subtle and progressive white matter (WM) alterations (Dayalu & Albin, 2015), have been observed early in HD progression, before the onset of motor symptoms (Aylward et al., 2011; Bourbon-Teles et al., 2019; Ciarmiello et al., 2006; De Paepe et al., 2019; Dumas et al., 2012; Faria et al., 2016; Gregory et al., 2018; McColgan et al., 2017; Paulsen et al., 2008; Ruocco et al., 2008; Shaffer et al., 2017; Tabrizi et al., 2009; Wu et al., 2017; Zhang et al., 2018).

An increasing body of research suggests that WM changes in HD are due to changes in myelin-associated biological processes at the cellular and molecular level (Bardile et al., 2019; Gómez-Tortosa et al., 2001; Huang et al., 2015; Jin et al., 2015; Myers et al., 1991; Radulescu et al., 2018; Teo et al., 2016; Yin et al., 2020) – for a critical review of such changes see Casella et al., (2020). Myelin, a multi-layered membrane sheath wrapping axons, is crucial for axonal structure and WM functionality (Martenson, 1992). In HD, myelin changes are suggested to follow both a topologically selective and temporally specific degeneration, with early myelinated fibres being the most susceptible to, and the first to be affected by, myelin breakdown (Bartzokis et al., 1999, 2007; Dumas et al., 2012; Faria et al., 2016; Phillips et al., 2013; Tabrizi et al., 2011).

The assessment of early myelin changes in the HD brain is therefore of fundamental importance for the understanding of disease pathogenesis and progression. Notably, as no disease-modifying treatment currently exists for HD, understanding the biological underpinning of HD-associated WM changes may prove useful for the identification of disease-related biomarkers, and for measuring responsiveness to pharmaceutical and other therapeutic approaches. Critically, given the certainty of onset in those that inherit the HD mutation, we can examine HD-associated myelin-related changes from the earliest, premanifest disease stages, with the potential to identify novel treatment targets for delaying disease onset.

### 1.2. Probing myelin changes in the HD brain with a multi-echo gradient-recalled echo (mGRE) sequence

Quantitative MRI of myelin affords valuable insight into myelin alterations and is thus of particular interest in the study of myelin-related disorders. Most neuroimaging studies that have quantified WM tissue properties in HD have used diffusion tensor magnetic resonance imaging (DT-MRI) (see Casella et al. 2020 for a review). However, while sensitive, DT-MRI measures are not specific to WM sub-compartments, challenging the interpretation of any observed change in these indices (Beaulieu, 2002; De Santis et al., 2014; Wheeler-Kingshott & Cercignani, 2009).

Other MRI techniques have the promise to provide much more myelin-specific information (MacKay & Laule, 2016). For example, myelin water imaging (MWI) quantifies the fraction of the faster decaying signal from water trapped between myelin lipid bilayers (MacKay et al., 1994), the so-called myelin water fraction (MWF). MWF has a good correlation with histological measurements of myelin, demonstrating its potential as an *in vivo* myelin marker (Laule et al., 2006; 2004; Webb et al., 2003). MWI techniques are typically based on spin-echo (MacKay et al., 1994) or mGRE sequences (Du et al., 2007). Interestingly, mGRE enables further characterisation of the myelin sheath by exploring its interaction with the magnetic field Bo, which is suggestively dependent on the g-ratio (i.e. the ratio of the inner-to-outer diameter of a myelinated axon) (Wharton & Bowtell, 2012).

A plethora of studies have demonstrated the non-mono-exponential nature of mGRE signal evolution with echo time (TE) in WM (e.g. Sati et al. 2013; Wharton and Bowtell 2012, 2013), arising from sub-voxel microstructure, with distinct signal components originating from water confined to the myelin, intra-axonal and extra-axonal water pools (Cronin et al., 2017; Nam et al., 2015b; Nunes et al., 2017; Sati et al., 2013; Tendler & Bowtell, 2019; Thapaliya et al., 2017; Wharton & Bowtell, 2012). As a result of the rapid T_2_* decay of the myelin water signal, the frequency of the total signal changes with TE, producing a local, microstructure-dependent contribution to the signal phase. However, in order to uncover the specific effects of microstructure on phase signal evolution, it is necessary to remove TE-dependent signal inhomogeneities resulting from non-local field variation, together with other non-TE-dependent phase changes, such as those due to radiofrequency interaction with the tissue (Schweser et al., 2011).

For this purpose, frequency difference mapping (FDM) has been presented recently as a phase-processing technique (Kleban et al., 2021; Sati et al., 2013; Schweser et al., 2011; Tendler & Bowtell, 2019; van Gelderen et al., 2012; Wharton & Bowtell, 2013). FDM is performed by comparing frequency maps acquired at short and long TEs so as to yield local frequency difference values which depend solely upon the underlying tissue microstructure, and in particular upon the local nerve fibre orientation with respect to the applied magnetic field. Critically, since both compartmentalization and myelination are prerequisites for the generation of frequency differences, FDM has great potential for the study of myelin changes in WM (Li et al., 2016; Wisnieff et al., 2015).

The aim of the present study was to exploit, for the first time in the HD literature, the sensitivity of the complex mGRE signal to WM microstructure, and particularly to myelin content, to assess callosal myelin changes at the premanifest stage of the disease. The corpus callosum (CC) is the largest WM fibre tract in the brain and carries information between the hemispheres; additionally, this tract plays an integral role in relaying sensory, motor and cognitive information between homologous cortical regions (Aboitiz et al., 1992), and provides vital connections to cortical areas known to be affected in HD (Crawford et al., 2013). Crucially, given its perpendicular orientation with respect to the B_0_ field, as a ‘proof of concept’ of the utility of the FDM approach in HD, the CC is a natural choice. Investigating this tract indeed afforded the largest possible frequency offsets in the myelin and axonal compartments, thus giving the most marked frequency difference ‘signature’ of myelin (Sati et al., 2013; Wharton & Bowtell, 2012; Yablonskiy et al., 2014).

Previous evidence from DT-MRI and volumetric studies has shown that changes in macro- and microstructure are detectable in the CC early in the disease course (Crawford et al., 2013; Di Paola et al., 2012, 2014; Phillips et al., 2013). The aim of the present work was to provide novel evidence on callosal changes in HD, and specifically to move beyond the existing literature by employing ultra-high field susceptibility measurements in order to afford a more biologically-meaningful interpretation of microstructure changes in this tract. Importantly, scanning participants at ultra-high field strength (i.e., 7T) afforded higher signal-to-noise ratio (SNR) and signal contrast (MacKay & Laule, 2016) per unit time, compared to more commonly-available field strengths (e.g. 3T).

Specifically, we sought to: i. establish the reliability of this method by investigating the anatomical variability in the reproducibility of FDM across the callosum at 7T; ii. compare two myelin-related parameters between premanifest HD patients and healthy controls; and iii. assess brain-function relationships in patients by exploring correlations between myelin content and cognitive function, as well as proximity to disease onset. The two myelin-sensitive metrics we assessed were: i. the myelin water signal fraction (f_m_) and ii. the difference in frequency offsets between myelin water pool and axonal water pool (Δω). The former is linked to the myelin volume fraction and may be used as a proxy for tissue myelin content (Laule et al., 2008; Li et al., 2015); the latter depends on the magnetic susceptibility difference and on the g-ratio, which is the ratio of the inner to outer diameter of myelinated axons (Wharton & Bowtell, 2012). Cognitive tests were selected in order to capture functioning across executive functions, working memory, social cognition and motor performance (Table 5), as these represent the earliest cognitive indicators of HD (Paulsen, 2011), and impaired performance in these domains has been associated with callosal microstructure changes (Kennedy & Raz, 2009; Lenzi et al., 2007; McDonald et al., 2018).

## 2. Materials and Methods

### 2.1. Subjects

#### Reproducibility study

To investigate the anatomical variability in the precision of the complex mGRE signal and the subsequent multi-compartmental analysis across the CC, six healthy subjects without known neurological or psychiatric conditions (3 female, 26-33 years-old) were scanned five times over a two-week period each. The study was approved by the Cardiff University School of Psychology Ethics Committee and written informed consent was obtained from all participants.

#### HD study

For the assessment of callosal myelin content in premanifest HD, MRI scans and cognitive tests were performed on 19 premanifest HD patients and 21 age, sex, and education matched healthy controls (Table 1). The study was performed with ethics approval by the local National Health Service (NHS) Research Ethics Committee (Wales REC 5 18/WA/0172); all participants provided written informed consent.

**Table 1.**
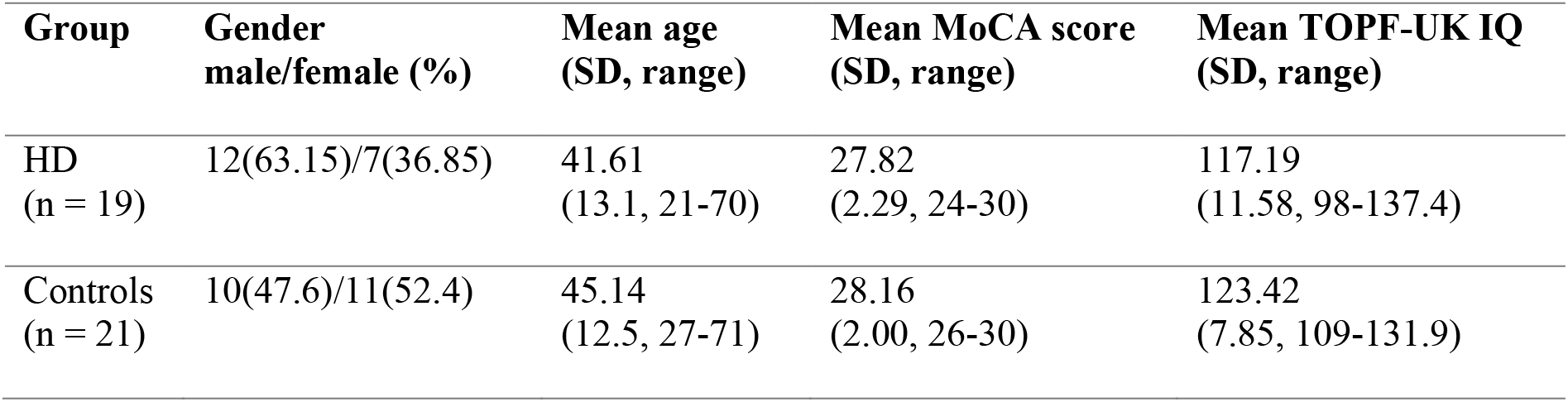
Summary of participants’ demographic and clinical background information. Age is displayed in years. MoCA = Montreal Cognitive Assessment out of 30; the higher the score the better the performance. TOPF-UKIQ = verbal IQ estimate based on the Test of Premorbid Functioning, UK version.

HD patients were recruited from the Cardiff HD Research and Management clinic, the Bristol Brain Centre at Southmead Hospital, and the Birmingham HD clinic at the Birmingham and Solihull NHS Trust. Healthy controls were recruited from Cardiff University, the School of Psychology community panel, and from patients’ spouses or family members.

In order to take part in the study, HD carriers had to be at the premanifest disease stage, and hence have no motor diagnosis, and to be enrolled in the EHDN Registry/ENROLL study (NCT01574053, https://enroll-hd.org). The progression of symptoms in ENROLL-HD participants is monitored longitudinally. As such, a full clinical dataset including medical history is available for each research participant, and some of these data were used for this study.

Table 1 summarizes information about demographic variables and performance in the Montreal Cognitive Assessment (MoCA) (Nasreddine et al., 2005) and in the Test of Premorbid Functioning - UK Version (TOPF-UK) for patients and controls. Although the two groups did not differ significantly in age, MoCA score, or TOPF-UK IQ, controls were on average slightly older and had a slightly higher IQ. Table 2 summarizes patients’ background clinical characteristics. Three individuals with CAG repeats of 37 (n = 1) and 38 (n = 2) were included in the current study. Although these individuals can be considered “affected”, they may have a lower risk of becoming symptomatic within their life span. Based on total motor scores (TMS), all patients were at the premanifest disease stage. Based on diagnostic confidence level scores (DCL), four patients presented some motor abnormalities, but none of them presented unequivocal motor signs of HD. Table 3 summarizes information about patients’ medication.

**Table 2.**
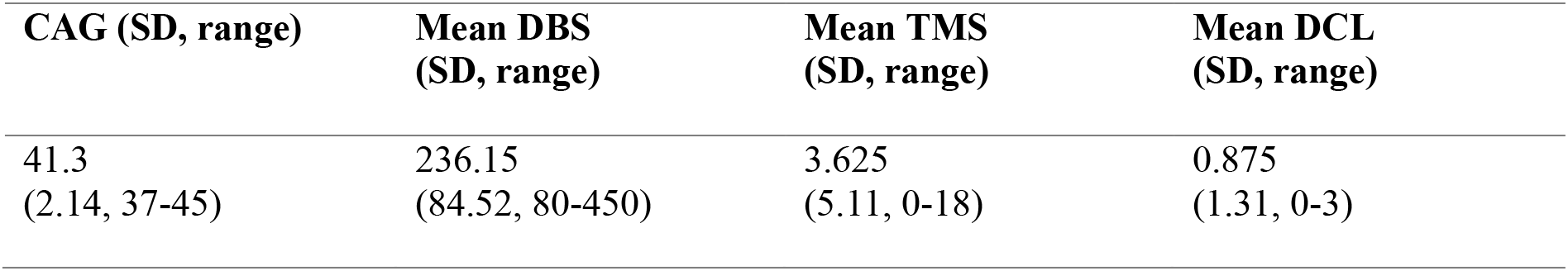
Background clinical information of the patients’ cohort. CAG = cytosine, adenine, and guanine repeat size. DBS = disease burden score, a measure of proximity to clinical onset of the disease (Tabrizi et al., 2012), calculated as follows: DBS = age × (CAG-35.5); the higher the DBS, the closer the patient’s proximity to disease onset. TMS = Total Motor Score out of 124 from the Unified Huntington’s Disease Rating Scale (UHDRS) Motor Diagnostic Confidence (Motor) – the higher the score stands for the higher the motor impairment. DCL = diagnostic confidence level, asks whether the participant “meets the operational definition of the unequivocal presence of an otherwise unexplained extrapyramidal movement disorder in a subject at risk for HD” (normal/no abnormalities = 0, nonspecific motor abnormalities = 1, motor abnormalities that may be signs of HD = 2, motor abnormalities that are likely signs of HD = 3, motor abnormalities that are unequivocal signs of HD = 4).

**Table 3.**
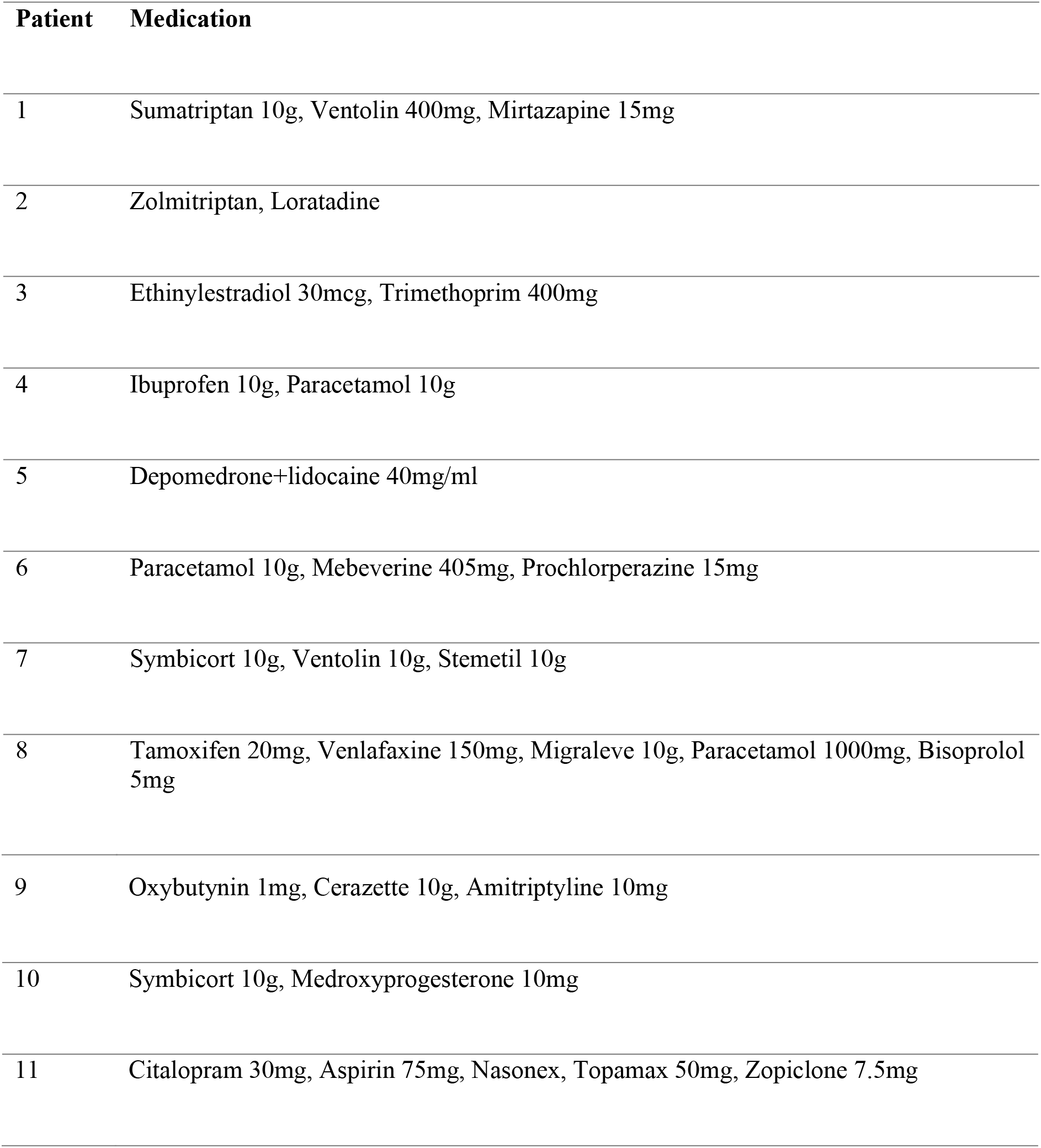
Information about patients’ medication. Out of the 19 patients we assessed, 11 had been on stable medication for four weeks prior to taking part in the study.

### 2.2. MRI data acquisition and processing

#### Imaging protocol

Complex multi-echo gradient-recalled echo (mGRE) data were acquired on a wholebody 7T research MR-system (Siemens Healthcare GmbH, Erlangen, Germany) equipped with a 32-channel head receive/volume transmit coil (Nova Medical).

By using a prototype mGRE sequence, a single mid-sagittal 5mm-thick slice was acquired, with in-plane field of view and resolution of 256×256 mm^2^ and 1×1 mm^2^, respectively. Acquiring a relatively thick slice afforded higher SNR and greater robustness in terms of slice misalignments across scans. The first echo time, echo spacing, and repetition time (TR) were set to TE1/ΔTE/TR = 1.62/1.23/100ms, the flip-angle of the radiofrequency excitation pulse was 15° and a total of 25 bipolar gradient echoes were acquired. Specifically, we acquired this 20 times, with read gradient polarities inverted half way through, with a total acquisition time of 8 minutes and 32 seconds.

#### Pre-processing steps

The complex data were reconstructed per receive channel, followed by a complex multiplication of signals acquired with opposite read-gradient-polarities, in order to remove phase shift between adjacent echoes (Kleban et al., 2021). Image-based coil-sensitivity-estimation was then used to perform coil combination (Bydder et al., 2002; Roemer et al., 1990). Frequency difference maps (FDM) were calculated from the phase data to correct for the RF-related phase offsets and the effects of the non-local B_0_ field inhomogeneities (Kleban et al., 2021; Tendler & Bowtell, 2019). Finally, a 3^rd^ degree spatial polynomial was fitted to the FDM data at each echo to correct for the residual eddy current effects. Figure 1 summarises the pre-processing steps. Additional details on pre-processing can be found in the Appendix.

**Figure 1.**
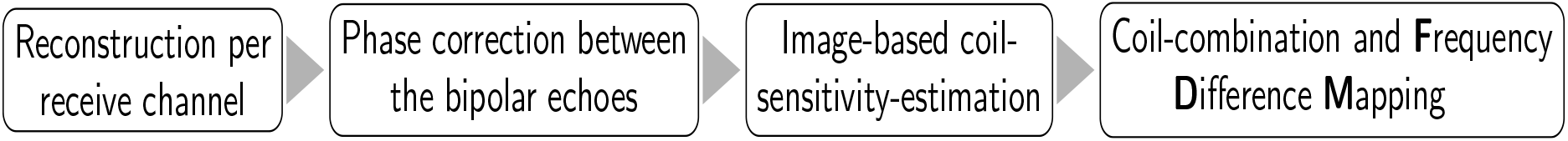
Schematic representation of the processing pipeline. FDM allowed removal of the RF-related phase offsets and linear effect of large-length-scale field perturbations, without perturbing the local non-mono-exponential WM signal.

#### Signal analysis

For each scan we manually segmented the corpus callosum from a magnitude mGRE image acquired at TE=15ms and further parcellated it into anterior, middle and posterior portions as shown in Figure 2. Magnitude and FDM data were averaged over each callosal segment.

**Figure 2.**
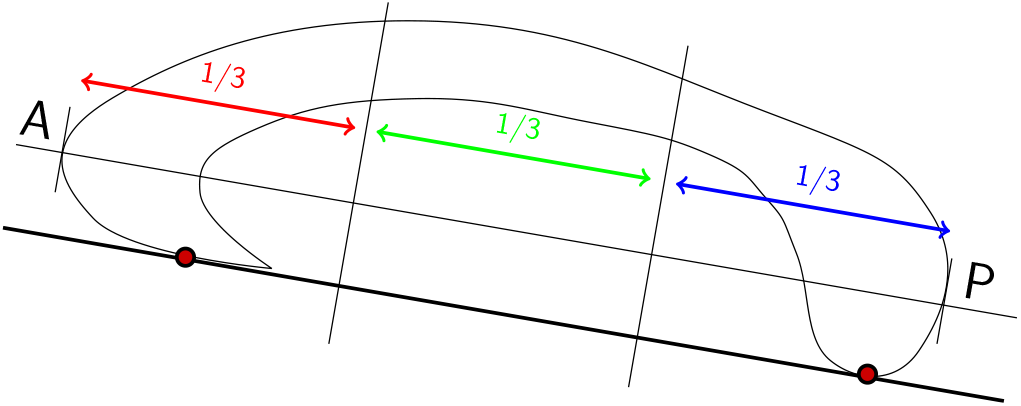
Schematic representation of the callosal segmentation protocol. The CC was segmented into three equal portions. Abbreviations: A = anterior; P = posterior.

FDM and magnitude signal evolution from each callosal segment were modelled using a three-pool-model of complex signal evolution (Figure 3a), where myelin water, intra-axonal and extra-axonal compartments each have different signal amplitudes, decay rates, and frequency offsets (Cronin et al., 2017; Nam et al., 2015b; Nunes et al., 2017; Sati et al., 2013; Tendler & Bowtell, 2019; Thapaliya et al., 2017; Wharton & Bowtell, 2013):

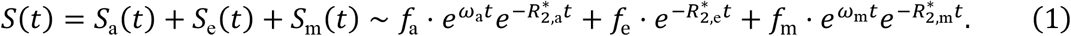

**Figure 3.**
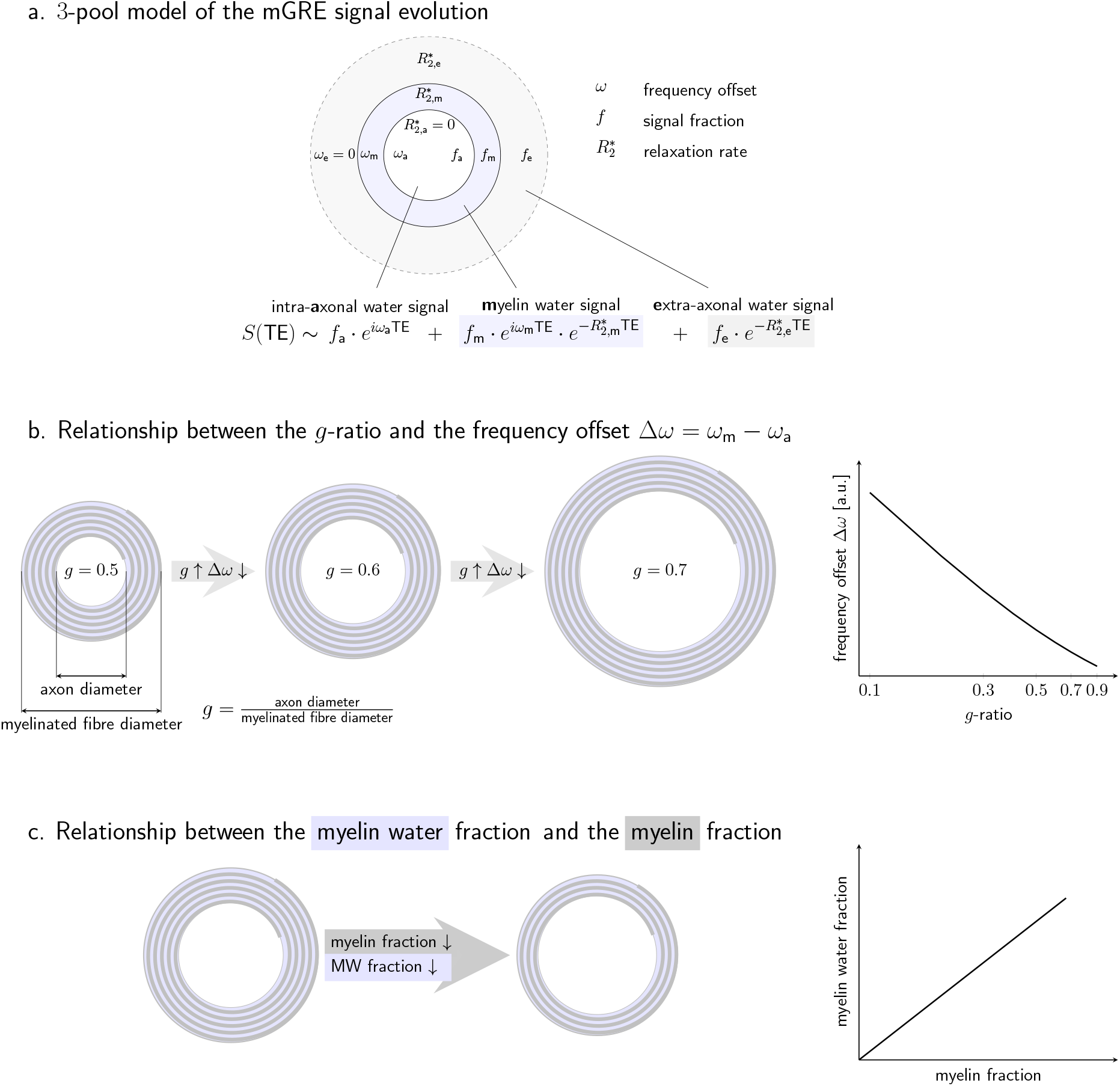
**a. Schematic representation of the three-pool model for describing mGRE signal evolution in WM fibres perpendicular to B_0_**. The signal is modelled as a superposition of complex myelin, intra- and extra-axonal water signals. **b. Schematic representation of the relationship between g-ratio and the frequency offset between myelin and axonal water pools.** An increase in g-ratio will be reflected by a decrease in Δω. **c. Schematic representation of the relationship between the myelin water fraction and the myelin fraction.** Such relationship highlights the potential of f_m_ as in vivo myelin marker.

Here, a, e, and m denote intra-, extra-axonal and myelin water and the complex mGRE signal *S*(*t*) is a superposition of their signals; *f* are the signal fractions, *ω* are the mean frequency offsets to the extra-axonal compartment, and 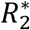 are the transverse relaxation rates.

By including the frequency offset characteristics of the different water compartments (Sati et al., 2013; Wharton & Bowtell, 2012), this model offers reliable myelin water estimation (Nam et al., 2015b; Sati et al., 2013; van Gelderen et al., 2012). Furthermore, previous research has suggested that ω_m_ and ω_a_ may both depend on the g-ratio (Wharton & Bowtell, 2012). We represent the dependence of the frequency offset Δω = ω_m_ – ω_a_ on the g-ratio in Figure 3b, while the relationship between the myelin water signal and myelin content is shown in Figure 3c.

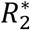-values of the slow-decaying intra-axonal signal were constrained to 0 because of relatively low maximum TE (31ms). This constraint helped to reduce the effect of the limited number of long TEs on the value estimation uncertainty at a cost of the potential under-/overestimation of the intra-/extra-axonal water signal fractions, respectively. Non-linear leastsquares fitting was performed with initial parameter estimates and fitting boundaries based on previous literature (Sati et al., 2013; Tendler & Bowtell, 2019; Thapaliya et al., 2017; Wharton & Bowtell, 2012, 2013) (Table 4).

Pre-processing and signal analysis were performed in Matlab (Matlab, The Mathworks, Natick, MA).

**Table 4.** Parameter values (initial and range) used in non-linear least squares fitting. Choice of fitting boundaries was based on previous literature (Sati et al., 2013; Tendler & Bowtell, 2019; Thapaliya et al., 2017; Wharton & Bowtell, 2012, 2013).

| Parameter | $f_m$ | $f_a$ | $f_e$ | $\omega_a/2\pi$<br>(Hz) | $\omega_m/2\pi$<br>(Hz) | $R_{2,a}^*$ (s<sup>-1</sup>) | $R_{2,m}^*$ (s<sup>-1</sup>) | $R_{2,e}^*$ (s<sup>-1</sup>) |
| --- | --- | --- | --- | --- | --- | --- | --- | --- |
| Initial value | 0.5 | 0.5 | 0.5 | -8 | 30 | 0 | 150 | 25 |
| Minimum value | 0 | 0 | 0 | -30 | 0 | 0 | 50 | 0 |
| Maximum value | 1 | 1 | 1 | 0 | 50 | 0 | 300 | 100 |

### 2.3. Cognitive tests

Cognitive performance was assessed in premanifest HD patients and age, sex and education matched healthy controls in the following tests: (1) the n-back task (Kirchner, 1958); (2) the digit span test from the Wechsler Adult Intelligence Scale-Revised (WAIS-R) (Wechsler, 1997); (3) the visual patterns test (Della Sala et al., 1997); (4) the Reading the Mind in the Eyes test **(Baron-Cohen et al., 2001)**, hereafter referred to as the eyes test; and (5) the finger tapping task (Freeman, 1940). Tests (3) and (4) were administered as paper and pencil tests, tests (1), (2), (5), and (6) by using the computerized version provided by the Psychology Experiment Building Language (PEBL) Test Battery (Mueller & Piper, 2014). Overall, we obtained a total of 6 cognitive outcome measures, which are summarised in Table 5, together with a short description of each task.

**Table 5.**
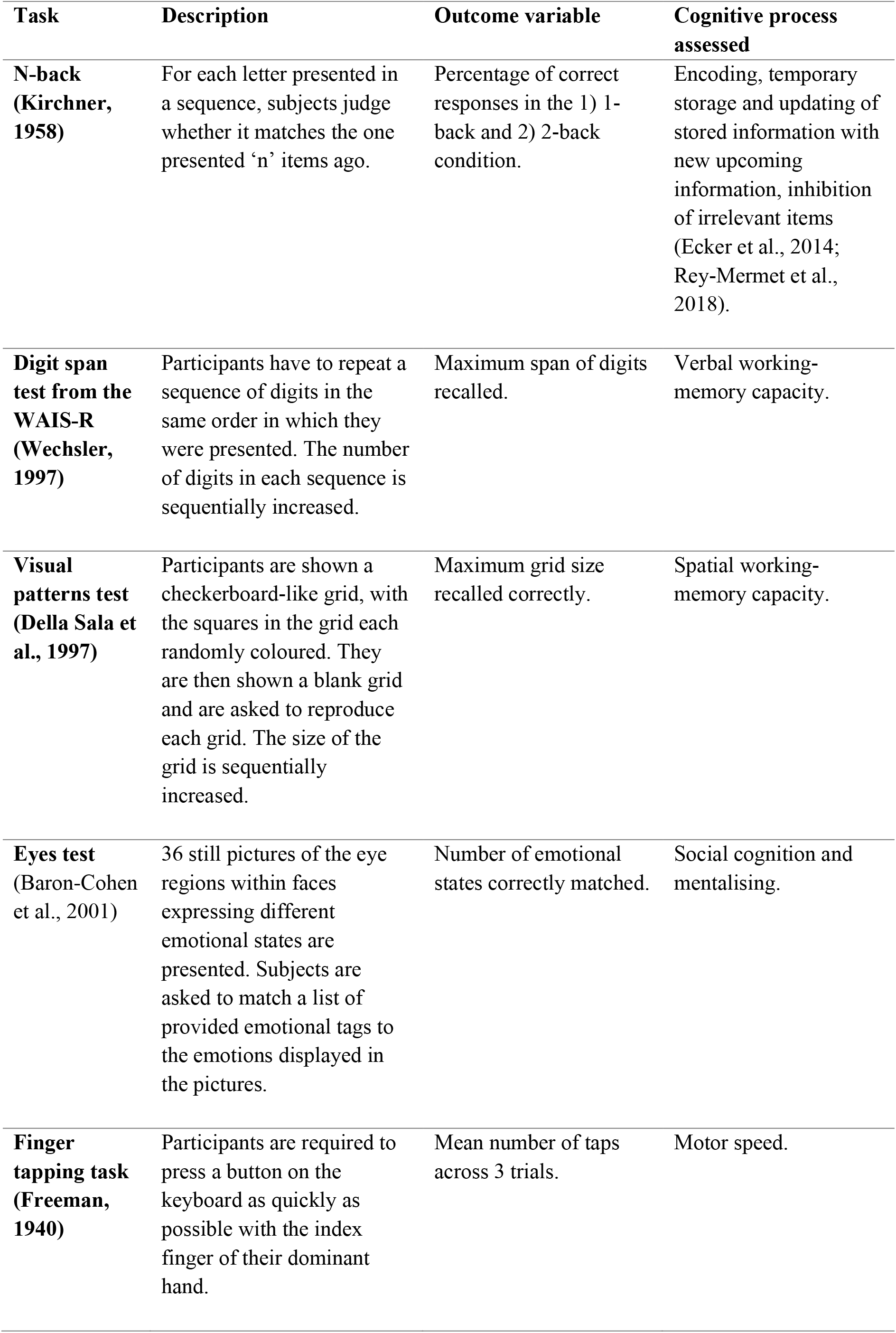
Cognitive outcome variables employed to assess patient-control differences in cognition. A short description of the task is provided, together with a list of outcome variables and cognitive domains assessed.

| **Task** | **Description** | **Outcome variable** | **Cognitive process assessed** |
| --- | --- | --- | --- |
| **N-back (Kirchner, 1958)** | For each letter presented in a sequence, subjects judge whether it matches the one presented 'n' items ago. | Percentage of correct responses in the 1) 1-back and 2) 2-back condition. | Encoding, temporary storage and updating of stored information with new upcoming information, inhibition of irrelevant items (Ecker et al., 2014; Rey-Mermet et al., 2018). |
| **Digit span test from the WAIS-R (Wechsler, 1997)** | Participants have to repeat a sequence of digits in the same order in which they were presented. The number of digits in each sequence is sequentially increased. | Maximum span of digits recalled. | Verbal working-memory capacity. |
| **Visual patterns test (Della Sala et al., 1997)** | Participants are shown a checkerboard-like grid, with the squares in the grid each randomly coloured. They are then shown a blank grid and are asked to reproduce each grid. The size of the grid is sequentially increased. | Maximum grid size recalled correctly. | Spatial working-memory capacity. |
| **Eyes test (Baron-Cohen et al., 2001)** | 36 still pictures of the eye regions within faces expressing different emotional states are presented. Subjects are asked to match a list of provided emotional tags to the emotions displayed in the pictures. | Number of emotional states correctly matched. | Social cognition and mentalising. |
| **Finger tapping task (Freeman, 1940)** | Participants are required to press a button on the keyboard as quickly as possible with the index finger of their dominant hand. | Mean number of taps across 3 trials. | Motor speed. |

### 2.4. Statistical analysis

We selected *f*_m_, ω_m_, ω_a_, and Δω = ω_m_ – ω_a_ obtained from mGRE complex signal analysis to perform further statistical analyses, as these metrics may reflect myelin changes in WM (see diagrams in Figure 3bc).

#### Reproducibility study

To assess the test-retest reproducibility of the data, we obtained the Fréchet distance (Fréchet, 1957) between FDM curves to measure their similarity. This method takes into account the location and ordering of points along the curves. Specifically, given two curves, Q and P, the Fréchet distance is defined as the minimum cord-length sufficient to join a point traveling forward along P and one traveling forward along Q. Furthermore, the coefficients of variation (CVs, the ratio of the standard deviation to the mean) across the 5 visits were computed for f_m_,, ω_a,m_, and Δω for each participant, for each segment. Finally, we used the R package cvequality (Version 0.1.3, Marwick and Krishnamoorthy 2019) to compute the ‘modified signed-likelihood ratio test for equality of CVs’ (Krishnamoorthy & Lee, 2014). This allowed us to test for significant differences between the CVs across the three segments, for each metric.

#### HD study

As greater measurement reproducibility was detected in the posterior segment of the CC, this was chosen as the region of interest for the assessment of patient-control differences in callosal myelin content. Age, but not TOPF-UK IQ, was found to be significantly correlated with both f_m_ and Δω. Hence age was included as a covariate in the analysis of group effects. Specifically, multiple regression analyses, assessing the effect of group, age, and a group-by-age interaction on f_m_ and Δω, were run in order to assess whether these metrics could disentangle age-related changes from pathologic HD-associated neurodegeneration. We performed regression diagnostics and examined QQ plots and outlier profiles to detect any values above or below the upper/lower boundary of 95% confidence intervals of the slope of the regression line. As a sanity check, we also confirmed our results by running robust linear regression analyses, using the *lmRob* R function from the robust package (Wang et al., 2009), which handles small sample sizes, skewed distributions and outliers (Wilkinson, 1999).

Principal component analysis (PCA) was employed to reduce the complexity of the cognitive data and hence the problem of multiple comparisons, as well as to increase experimental power. We examined potential confounding effects of age or TOPF-UK IQ on the extracted components. Age was included as a covariate in the model, which explored the effect of group, age, and a group-by-age interaction on the scores on the extracted components. We interpreted any significant effect by referring to the variables with significant component loadings, as highlighted in bold in Table 7.

As a significant group effect was detected on f_m_, Spearman’s rho correlation coefficients were calculated in patients between this metric, DBS, and scores on the first cognitive component (i.e., the component capturing the greatest amount of variability in the data), to assess disease-related brain-function relationships. We corrected for multiple comparisons with the Bonferroni correction with a family-wise alpha level of 5% (two-tailed). Significant correlations were further assessed with partial correlations to control for age as potentially mediating variable.

All statistical analyses were carried out in R Statistical Software (Foundation for Statistical Computing, Vienna, Austria) and Matlab (Matlab, The Mathworks, Natick, MA).

## 3. Results

### 3.1. The precision of FDM in the corpus callosum is anatomically variable

Figure 4 shows an example of FDM and magnitude signal evolution as a function of TE, for anterior/middle/posterior callosal segments, and the corresponding images at TE=15ms for five visits.

**Figure 4.**
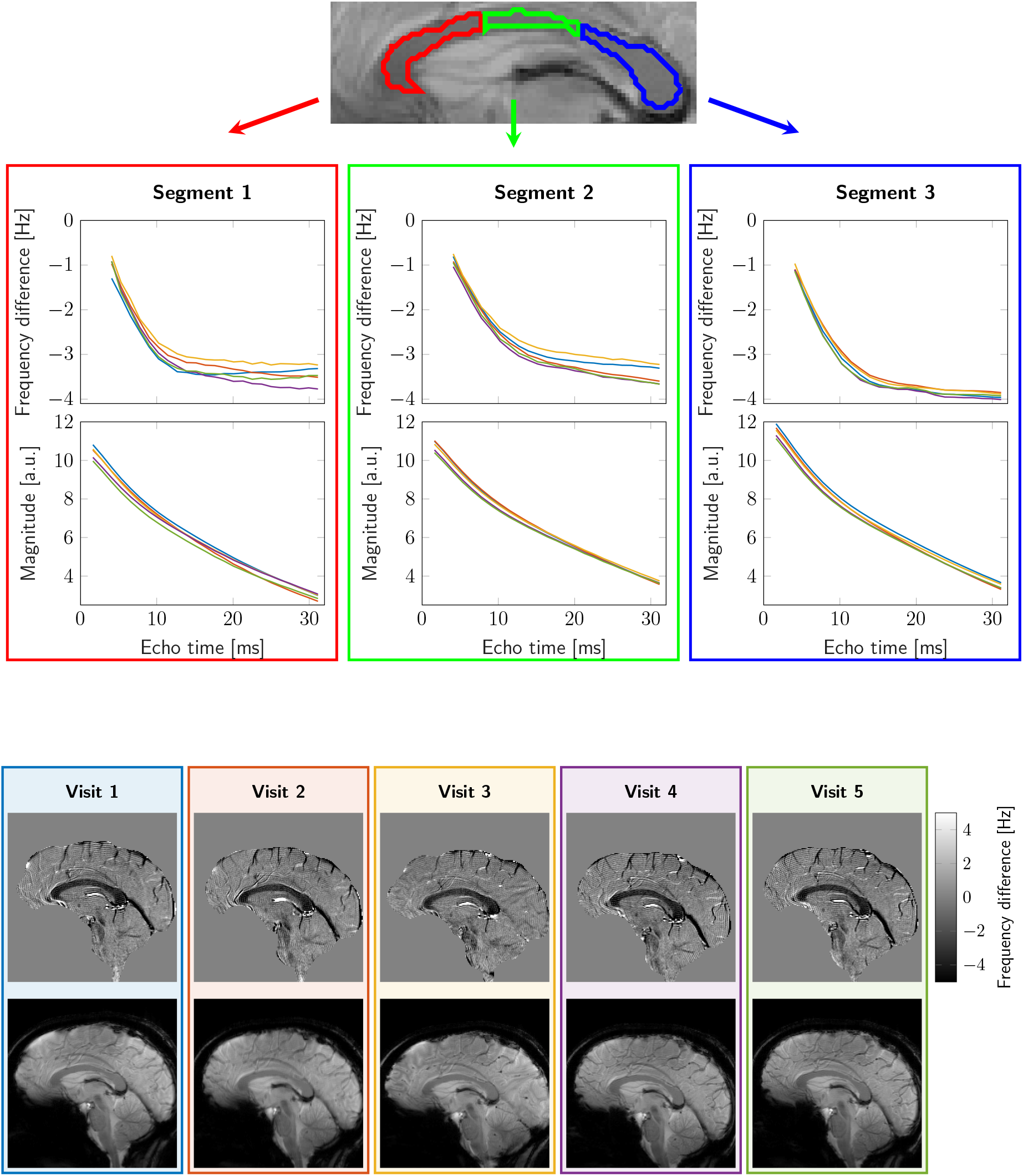
Example reproducibility data from one representative subject. The top image shows the callosal parcellation overlaid on a magnitude image; the plots show frequency difference and magnitude of signal as function of TE for anterior/middle/posterior callosal segments; corresponding frequency difference and magnitude images at TE=15ms for five visits are shown at the bottom.

Analysis of curve similarity (Figure 5) and fitting parameter repeatability (Figure 6, Table 6) suggested greater reproducibility of measures in the posterior callosal portion as compared to the anterior sections. This is shown by generally lower Fréchet distance values and by overall low coefficients of variation calculated for the fitting parameters, which ranged between 3.72% and 12.02%, in this region (Table 6). The modified signed-likelihood ratio test for equality of CVs confirmed that CVs in the posterior callosal segment were significantly smaller across all measures [f_m_: p = 0.02; ω_a_/2π (Hz): p < 0.001; ω_m_/2π (Hz): p < 0.001; Δω//2π) (Hz): p = 0.020].

**Figure 5.**
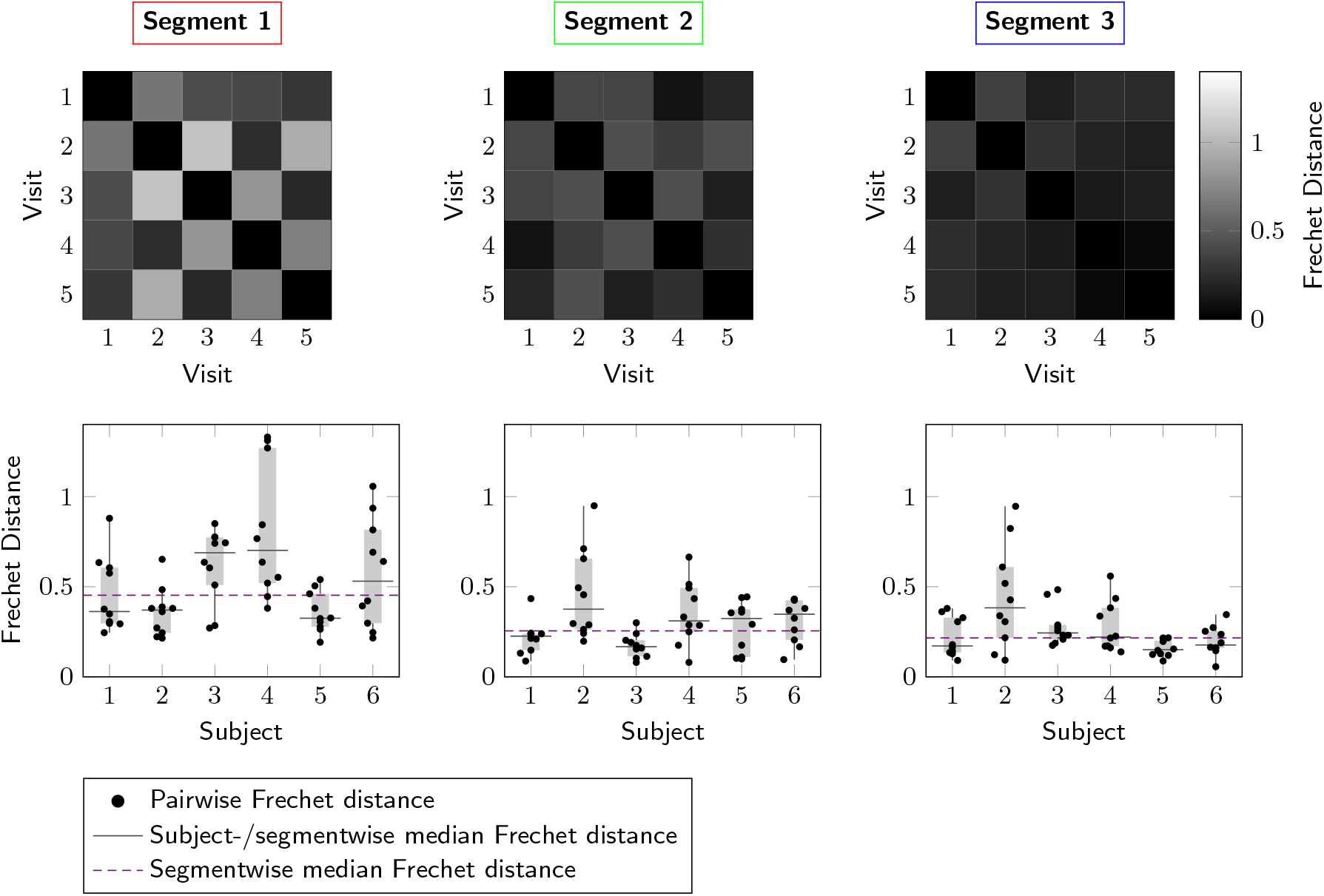
Assessment of the similarity of frequency difference curves. **Top:** Fréchet distance matrices from a representative participant for the three callosal segments. **Bottom**: Repeatability of frequency difference evolution across five visits for all subjects.

**Figure 6.**
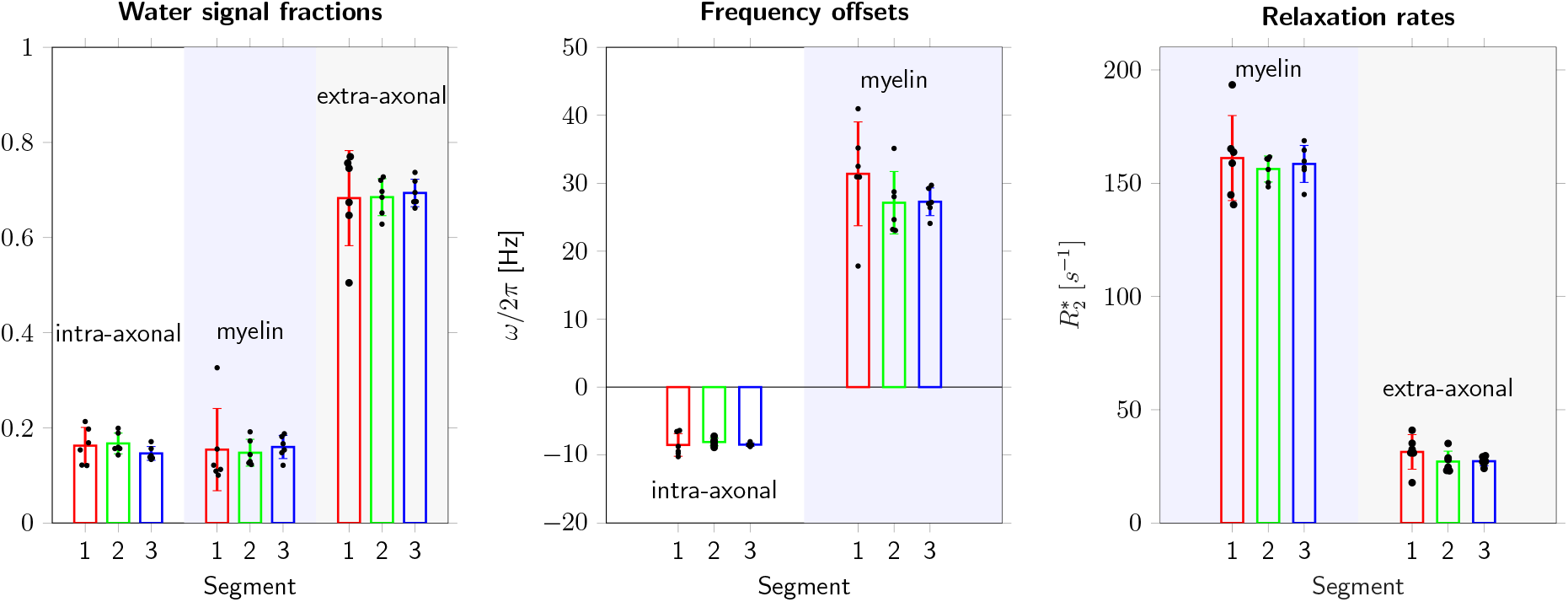
Fitting parameters estimated from the three-pool model, grouped for all subjects. Colours represent the three callosal segments (red/green/blue for anterior/mid/posterior segments, respectively). The first plot shows signal fractions of myelin, intra-/extra-axonal water; the frequency offsets are displayed in the middle plot; the last plot shows the relation rates of myelin and extra-axonal water signals.

**Table 6.**
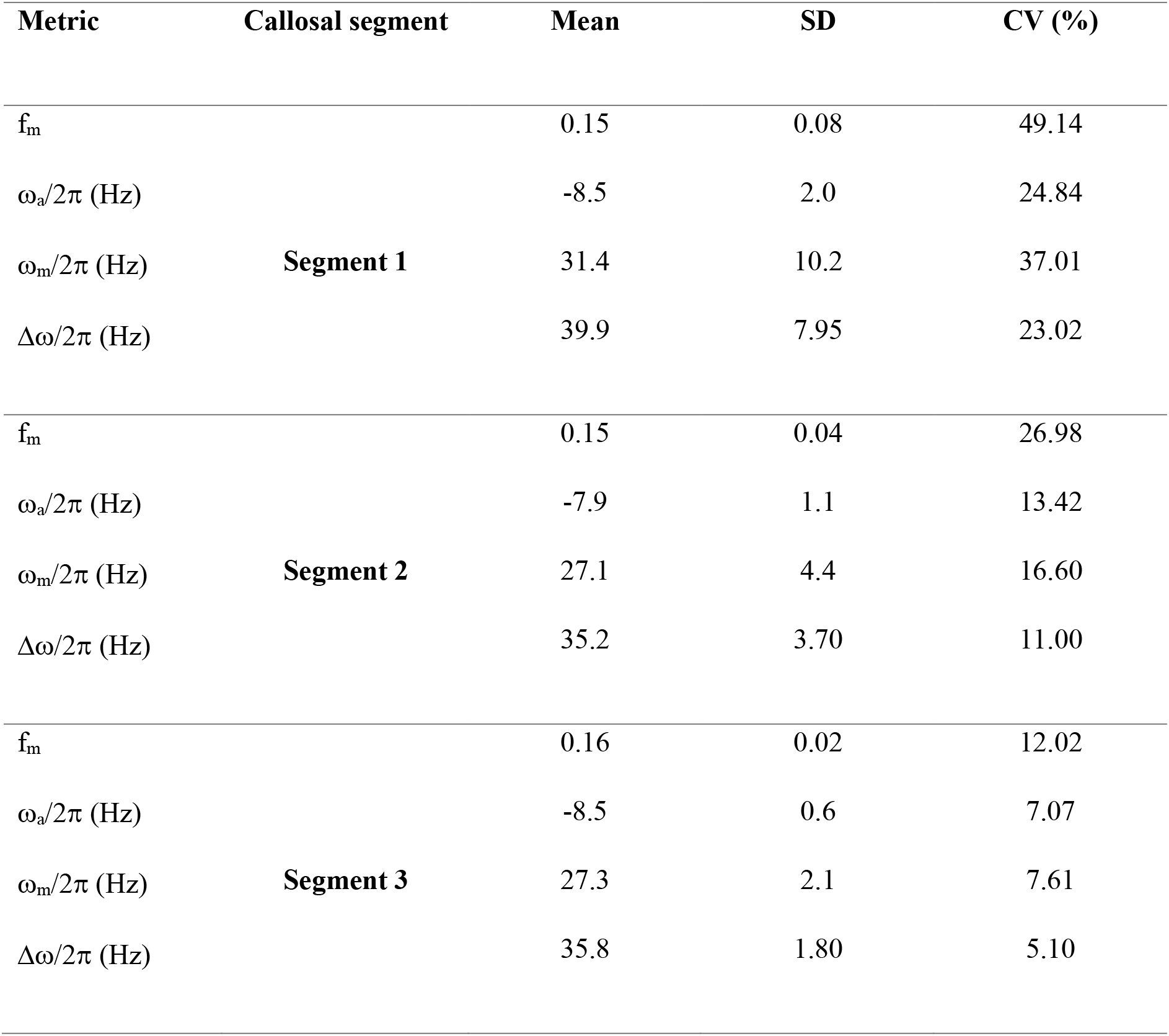
Reproducibility of myelin water signal fraction (f_m_), frequency offsets of axonal (ω_a_/2π) and myelin (ω_m_/2π) water pools, and difference in frequency offsets between myelin and axonal water pools (Δω/2π)). Means, standard deviation (SD) and coefficients of variation (CV) are reported for each value, across the different segments.

| Metric | Callosal segment | Mean | SD | CV (%) |
| --- | --- | --- | --- | --- |
| $f_m$ | **Segment 1** | 0.15 | 0.08 | 49.14 |
| $\omega_a/2\pi$ (Hz) | | -8.5 | 2.0 | 24.84 |
| $\omega_m/2\pi$ (Hz) | | 31.4 | 10.2 | 37.01 |
| $\Delta\omega/2\pi$ (Hz) | | 39.9 | 7.95 | 23.02 |
| $f_m$ | **Segment 2** | 0.15 | 0.04 | 26.98 |
| $\omega_a/2\pi$ (Hz) | | -7.9 | 1.1 | 13.42 |
| $\omega_m/2\pi$ (Hz) | | 27.1 | 4.4 | 16.60 |
| $\Delta\omega/2\pi$ (Hz) | | 35.2 | 3.70 | 11.00 |
| $f_m$ | **Segment 3** | 0.16 | 0.02 | 12.02 |
| $\omega_a/2\pi$ (Hz) | | -8.5 | 0.6 | 7.07 |
| $\omega_m/2\pi$ (Hz) | | 27.3 | 2.1 | 7.61 |
| $\Delta\omega/2\pi$ (Hz) | | 35.8 | 1.80 | 5.10 |

### 3.2. Multi-compartment analysis reveals HD-related myelin changes in the posterior segment of the CC

Figure 7 plots the relationship between age and f_m_, and between age and Δω, split by group, in Segment 3 of the CC. Group and age explained 45% of the variance in f_m_ [R^2^ = 0.45, F(4, 35) = 7.18, p < 0.001]. Specifically, it was found that group significantly predicted f_m_ values in this portion of the CC (β = −0.13, p = 0.03), as did age (β = −0.82, p < 0.001). However, we did not detect a significant group-by-age interaction effect (β = 0.82, p = 0.08) (Figure 7, left). Overall, HD patients presented a flatter age-related variation in this metric, with values being overall lower, especially in younger subjects.

**Figure 7.**
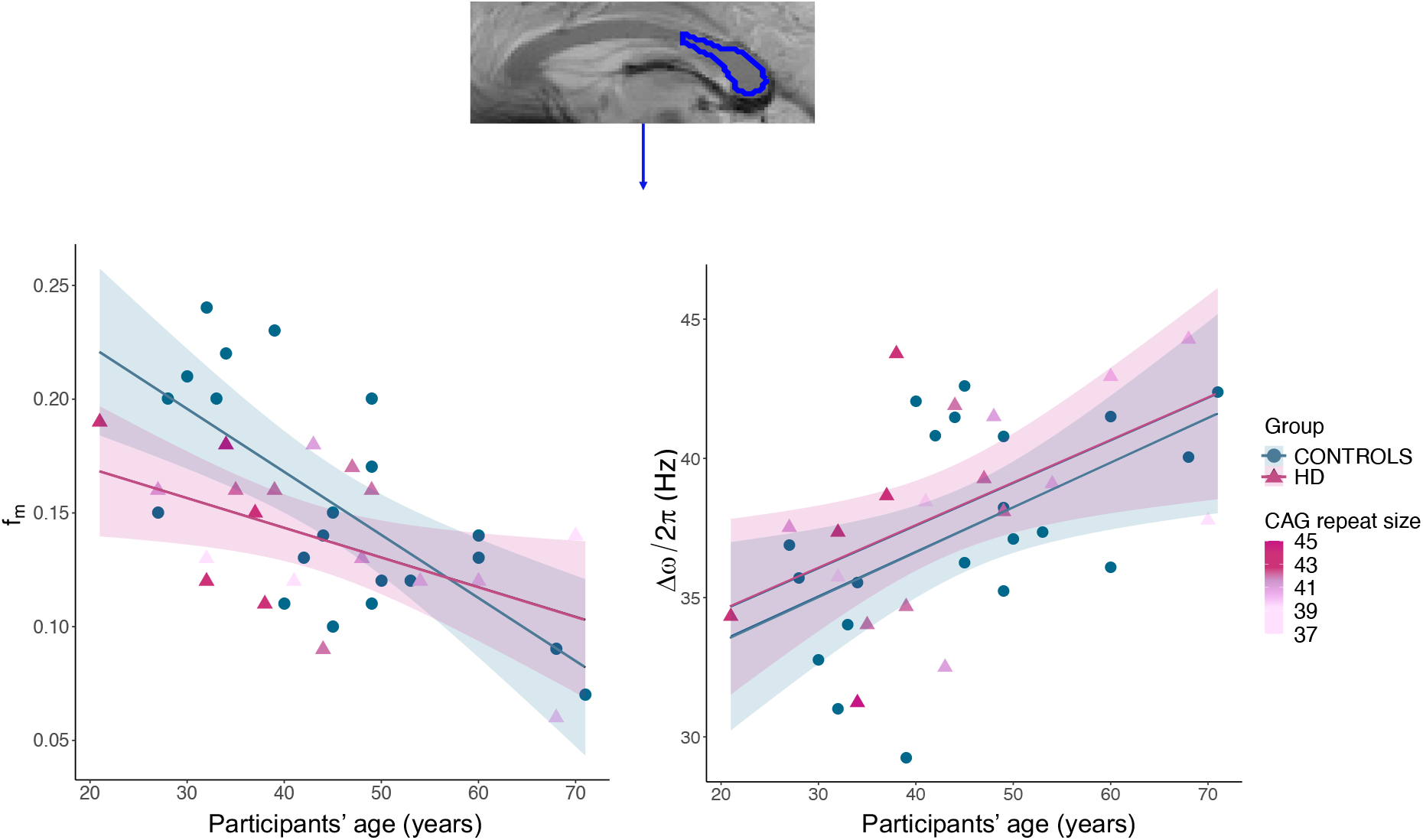
**Top: Parcellation of segment 3 of the CC overlaid on a magnitude image**; the same protocol as the one detailed in the reproducibility study section was utilised. **Left: Regression plot showing the relationship between age and f_m_. split by group.** Both age and group were significant predictors of variance in f_m_. No significant interaction effect was detected. **Right: Regression plot showing the relationship between age and Δω, split by group.** Age was a significant predictor in the model, while group did not significantly predict variance in this metric. No significant interaction effect was detected. HD data points are coloured by CAG repeat size: older HD carriers presented shorter CAG repeat mutation and a trend for a greater overlap in f_m_with values of age-matched healthy controls, indicating that CAG repeat size may directly affect myelin content in premanifest HD.

On the other hand, age and group explained 31% of the variance in Δω [R^2^ = 0.31, F(4, 35) = 3.87, p = 0.01]. Age was found to be a significant predictor of variance in this metric (β = 0.55, p = 0.009), so that being older was associated with a greater Δω in this sample. However, belonging to the patient or the control group did not have a significant effect on this measure (β = 0.13, p = 0.8) (Figure 7, right).

### 3.3. Premanifest HD is associated with greater age-related decline in executive functions

With PCA of cognitive test scores, the first three components explained 77.7% of the variability in performance in the administered tests (Figure 8). The first component loaded positively on variables from the n-back task and was therefore summarized as an “executive function/updating” component. Principal component 2 (PC2) was summarised as a “visuo-spatial motor function” component. Finally, PC3 was summarized as a “verbal working memory capacity” component as this loaded mostly on the digit span task.

**Figure 8.**
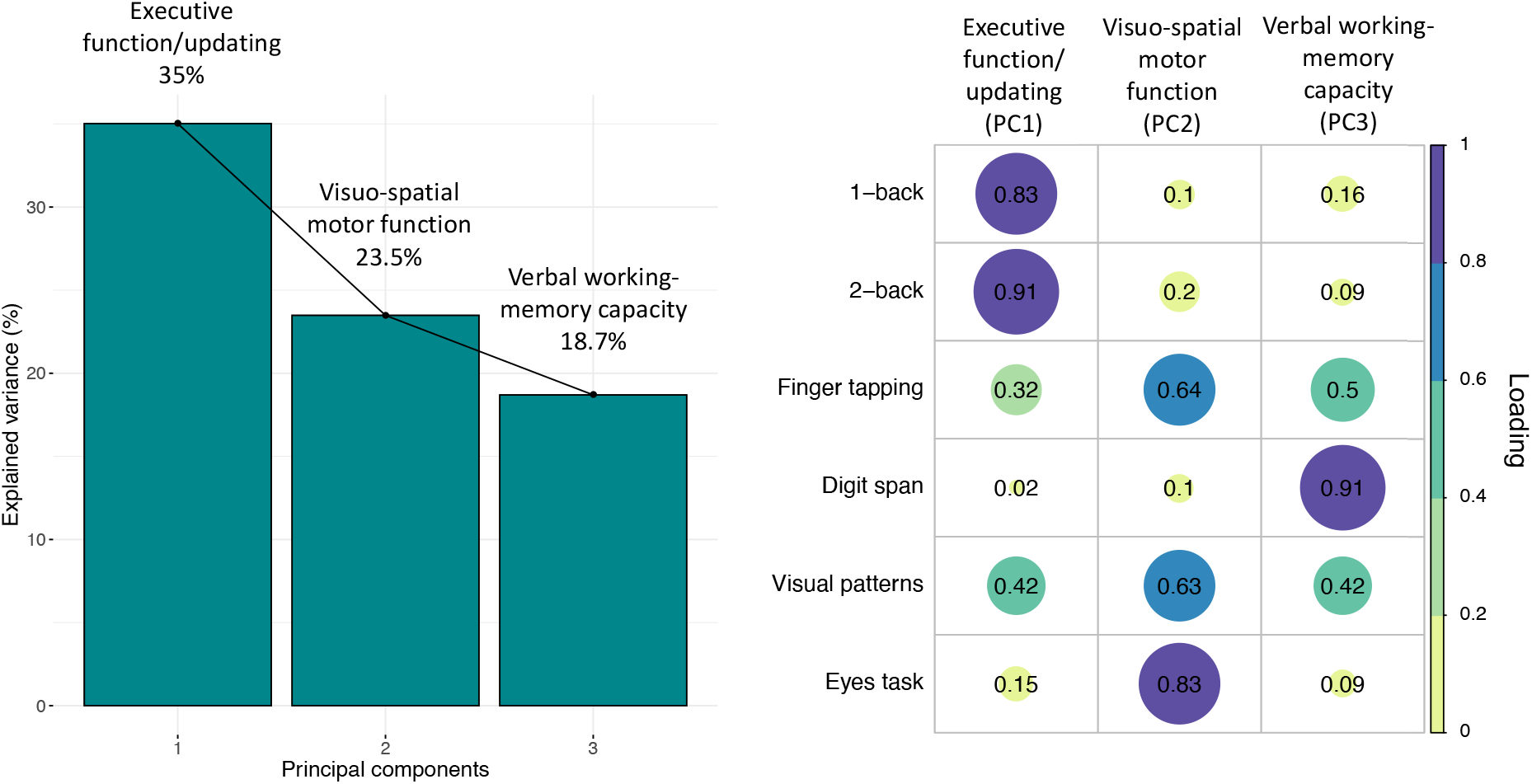
PCA of the cognitive data with varimax rotation. **Left: PCA scree plot. Right: Plot summarising how each variable is accounted for in every principal component.** The absolute correlation coefficient is plotted. Colour intensity and the size of the circles are proportional to the loading. Three components explaining over 77% of the data variability were extracted. PC1 loaded on n-back task performance and was therefore summarized as “executive function” component; PC2 was summarized as “visuo-spatial motor function” component; PC3 loaded on digit span task performance and was therefore summarized as “working memory” component. 7 control cases did not complete all tests and were therefore excluded from the PCA. The final sample size for the PCA was n=19 for the HD group and n=14 for the control group.

We detected no significant main effect of group (although a trend was present) (β = 1.9, p = 0.059) or age (β = 0.004, p = 0.815) on the executive function/updating component, but a significant interaction effect was present between group and age [β = −0.06, p = 0.006, R^2^ = 0.52, F(3,28) = 10.6, p = 0.006], indicating that while younger HD patients present executive function scores which tend to overlap with those of healthy controls, the gap in performance between the two groups is significantly larger at later ages (Figure 9, left).

On the other hand, the main effects of group (β = −1.71, p = 0.23) and age (β = −0.036, p = 0.14), and the interaction effect between group and age (β = 0.03, p = 0.31) on the visuo-spatial motor function component were not significant [R^2^ = 0.09, F(3,28) = 0.94, p = 0.43]. Similarly, we did not detect a significant main effect of group (β = 0.03, p = 0.82) and age (β = 0.03, p = 0.15), nor a significant interaction effect between group and age (β = −0.003, p = 0.91) on the verbal working memory capacity component [R^2^ = 0.14, F(3,28) = 1.63, p = 0.20].

### 3.4. Interindividual differences in myelin content in the posterior callosum relate to executive function but not to proximity to disease onset in premanifest HD patients

We detected a significant positive correlation between the patients’ inter-individual variation in f_m_ and their scores on the executive component (r = 0.542, p = 0.02, corrected p = 0.04) (Figure 9, right); although a positive trend remained, this relationship was no longer significant after partialling out age (r = 0.37, p = 0.13). Additionally, there was no association between inter-individual variation in f_m_ and proximity to disease onset as measured with DBS (r = −0.09, p = 0.686, corrected p = 1).

**Figure 9.**
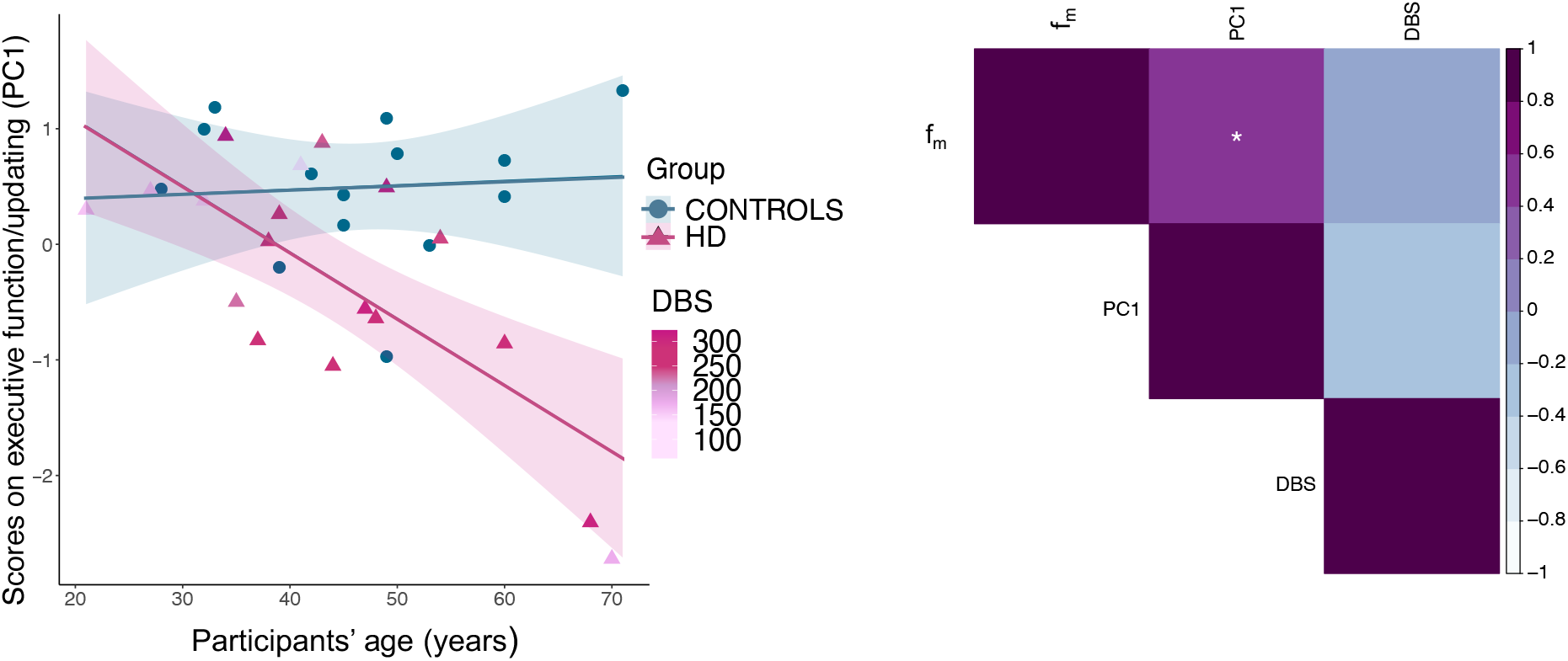
**Left: Relationship between executive function scores and age, in patients and controls**. A significant interaction effect between group and age was detected, suggesting that the group difference in executive function scores is larger at later ages. HD data points are coloured by DBS. Older HD patients tend to be closer to disease onset, possibly confounding the effect of age on this measure. **Right: Relationship between f_m_ executive function scores and DBS in patients, Bonferroni-corrected.** A significant positive correlation was found between f_m_ and executive function scores. Colour intensity is proportional to the strength and direction of the correlation. * p < 0.05, ** p < 0.01, *** p < 0.001, Bonferroni-corrected.

## 4. Discussion

### 4.1. Reproducibility of the mGRE signal across the CC

The reproducibility of FDM evolution and 3-pool model derivatives in the CC was found vary according to anatomical location, with more precise fitting parameters in the posterior portion of the callosum. Lower reproducibility of the data in the anterior portions of the CC could be attributed to in-flow artifacts from the anterior cerebral artery (Nam et al., 2015a; Tendler & Bowtell, 2019). Potential solutions and their limitations have been discussed in previous work, such as the application of flow saturation RF pulses to the inferior portion of the head (Nam et al., 2015a).

The estimated fitting parameters for the intra-axonal and myelin water frequency offsets (−7.9 to −8.5 Hz and 27.1 to 37.4 Hz, respectively) and the myelin water signal fraction (0.15 to 0.16) were consistent with previously reported values (Sati et al., 2013; Tendler & Bowtell, 2019; Thapaliya et al., 2017; Wharton & Bowtell, 2012). On the other hand, the relative signal fraction between intra- and extra-axonal compartments was lower than in other studies (Tendler & Bowtell, 2019). This could be attributed to the constraint we placed on the intra-axonal water 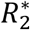 when modelling the data, which we introduced in order to reduce the effect of the limited number of long TEs on the value estimation uncertainty. Please refer to Appendix B for simulations we conducted on the impact of setting 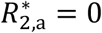 on MW signal fraction values. Finally, slice positioning relative to the CC, and the subsequent ROI segmentation, may also have introduced some variability in test-retest scans signal and estimated parameters.

### 4.2. Application of the 3-pool model to assess myelin content in premanifest HD

We observed significantly lower f_m_ values in premanifest HD patients compared to healthy controls, suggesting the presence of myelin impairment (Bartzokis et al., 2007). Previous animal studies have already shown a link between HD pathology and changes in myelin-associated biological processes at the cellular and molecular level (Bardile et al., 2019; Huang et al., 2015; Jin et al., 2015; Teo et al., 2016). Crucially, however, previous *in vivo* investigations of WM changes in HD have predominantly employed indices from DT-MRI (Pierpaoli & Basser, 1996) or magnetization transfer ratio (MTR) imaging (Henkelman et al., 2001). These indices may be influenced by a multitude of processes affecting tissue microstructure and biochemistry (Beaulieu, 2002; De Santis et al., 2014; Harsan et al., 2006; Henkelman et al., 1993; Wheeler-Kingshott & Cercignani, 2009). In contrast, our study exploited FDM and a three-pool modelling of the mGRE signal to afford improved WM compartmental specificity, and to estimate f_m_ as a marker of myelin content. Importantly, histological evidence shows that this metric is less sensitive to concomitant pathological processes such as inflammation (Gareau et al., 2000), suggesting that this may be a more specific measure of tissue myelination than other MRI-derived measures. Our results highlight the potential of f_m_ in helping to better understand HD pathogenesis and progression, and to gain further insight into the biological basis of WM microstructural changes in the HD brain. Additionally, they suggest the presence of myelin breakdown as an early feature of HD progression and are consistent with evidence from a quantitative magnetisation transfer study (Bourbon-Teles et al., 2019), which demonstrated reductions in the macromolecular proton fraction - a myelin sensitive measure - in HD patients.

In this study, we did not detect a group effect on Δω, while this parameter was shown to significantly increase with age. Based on the model proposed by Wharton and Bowtell (2012), this might reflect two processes: i. the g-ratio decreases with age; ii. the magnetic susceptibility difference between the myelin sheath and the extra-axonal pool increases with age. The first suggestion contradicts previous studies showing g-ratio increases with age (Peters, 2009). However, as fibres with smaller diameters tend to have slightly lower g-ratios (Berthold et al., 1983), a selective loss of large-diameter axons would lead to an overall reduced voxel-averaged g-ratio (Cercignani et al., 2017). This scenario is a plausible explanation of our finding, as fibres in the posterior portion of the CC are predominantly large and early myelinated (Aboitiz et al., 1992). Additionally, an increase in iron-containing glial cells in the surrounding extra-axonal space could result in the increase of the susceptibility difference between myelin sheath and the extra-axonal space (Xu et al., 2015). This is consistent with evidence showing that over the course of aging, iron accumulates in the brain (Connor et al., 1990; Dexter et al., 1991; Jellinger et al., 1990; Zecca et al., 2004).

With regards to the cognitive assessments, our results indicate that executive functions, and specifically the updating of relevant information, tend to deteriorate to a larger extent with age in HD patients, compared to controls. Nevertheless, these results might have been confounded by stage of disease progression, as older participants presented a higher DBS. Understanding of the nature of cognitive deficits associated with HD pathogenesis and progression provide useful guidance for future research into the efficacy of cognitive training and rehabilitation approaches in HD (Andrews et al., 2015). Our findings suggest that such approaches might be more effective early in the lifetime and in disease progression.

In the present study, we also found that patients’ inter-individual variability in f_m_ was positively associated with their scores on the executive/updating component. Although anterior, rather than posterior, callosal portions have been normally associated with frontal-lobe-mediated executive functions (e.g. Jokinen et al. 2007), posterior callosal fibres are connected to posterior parietal areas of the brain (Goldstein et al., 2020); these areas have been associated with top-down modulation during inhibition and attention processes (Erickson et al., 2009; Hopfinger et al., 2000), which are recruited during maintenance and updating of relevant information. It is therefore plausible that microstructural variation in this callosal segment may impact performance in this cognitive domain. Additionally, although a positive trend remained, this relationship was no longer significant after partialling out age, stressing the important role of aging in both myelin content in the brain, and executive functioning (Grieve et al., 2007; Guttmann et al., 1998; Lintl & Braak, 1983; Pakkenberg et al., 2003). Interestingly, we found no association between inter-individual variation in f_m_ and proximity to disease onset as measured with DBS. This suggests that myelin differences may precede the onset of clinical symptoms in HD and may not directly relate to disease stages.

### 4.3. Challenges to the interpretation of mGRE signal evolution

In this work, the complex WM mGRE signal was modelled as a superposition of three water signals which were assigned to the MW, intra-, and extra-axonal pools. Similar to previous work, the pool with the shortest 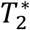 and the largest (positive) frequency offset was assigned to the myelin water (MW) pool. This representation assumes that chemical exchange between the MW pool and the intra-/extra-axonal water pools is negligibly small and that the myelin sheath is the dominant source of magnetic susceptibility effects in WM.

However, it has previously been suggested that magnetisation exchange can affect the interpretation of tissue compartmentalisation from MR signals or NMR frequency in brain tissues (Shmueli et al., 2011; van Gelderen & Duyn, 2019). Other work has suggested that the presence of iron-rich oligodendrocytes and the geometry of axons may have an impact on compartmental frequency distributions and consequently on the MR signal (Xu et al., 2015, 2018). Additionally, a previous study performed MWI using *T*_2_-weighted techniques on a *postmortem* brain before and after iron extraction by reductive dissolution and reported a 26% reduction in the MW signal fraction (Birkl et al., 2019). Based on the reportedly excellent agreement between mGRE and *T*_2_-weighted MWI techniques at 3T (Alonso-Ortiz et al., 2018b), it is plausible that iron deposition may have altered the MW signal fraction in our mGRE experiments at 7T. Finally, the similarity between intra- and extra-axonal mGRE signal evolutions may further confound the accuracy of compartmentalisation, especially at lower magnetic fields and short echo times (Chan & Marques, 2020). Crucially, this work exploited ultra-high magnetic fields and this may have allowed a better separation between the intra- and extra-axonal signals (Alonso-Ortiz et al., 2018a). Nevertheless, simultaneous variation of multiple experimental parameters and additional information derived from other MRI contrasts may further improve mGRE signal interpretation (Chan & Marques, 2020; Kleban et al., 2020).

## 5. Conclusions and future directions

In summary the present study exploited, for the first time in HD research, the sensitivity of the 3-pool model analysis of the complex mGRE signal to quantify myelin changes in premanifest HD. Results stress the potential of this marker in helping to better understand HD pathogenesis and progression, and provide original in-vivo evidence for reductions in f_m_, a proxy MRI marker of myelin, in human premanifest HD. Expanding on evidence from pathology and animal studies, our results suggest that myelin breakdown is an early feature of HD progression, and lend supporting evidence to the progression model suggesting that early myelinated fibres are affected by myelin breakdown early in the disease (Bartzokis et al., 2007).

Our findings were based on a relatively small sample size and warrant replication in larger samples. In addition, three individuals with CAG repeats of 37 and 38 were included in the current study. Though these individuals can be considered “affected”, they may have a lower risk of becoming symptomatic within their life span, and some studies have chosen to exclude individuals with reduced disease penetrance (e.g. Rubinsztein et al., 1996). Although the aim of this study was to look at the premanifest, rather than symptomatic, stage of the disease, their inclusion in the current study may raise the possibility of Type II errors when generalizing to the wider population of individuals with HD. Though this did not seem to be the case in the present study as we did detect a group effect, future studies may want to replicate these results in a sample of premanifest HD patients with full disease penetrance. Finally, while it is tempting to assign, unequivocally, a one-to-one correlation between changes in MRI signal and biological properties, and thus interpret these changes purely in terms of changes in myelination, these findings need to be interpreted with caution. For example, changes in iron content cannot be ruled out.

Future studies should assess HD-related changes in f_m_ longitudinally rather than cross-sectionally and investigate how these changes relate to clinical symptoms over time, to further understand the utility of this metric as a marker of early disease development and progression. Additionally, this study utilised a single-slice technique, and investigated a small portion of the CC, thus limiting the assessment of global diffuse tissue damage. Of special interest for future investigations might be to increase brain coverage and assess how f_m_ changes may differentially impact early and later myelinating regions in the premanifest HD brain. With regards to the observed increase of Δω with age, MR axon radius mapping using diffusion MRI and ultrastrong gradients (Veraart et al., 2020, 2021) may help elucidate whether a selective loss of large-diameter axons is producing this effect.

## Abbreviations

CAG: cytosine, adenine, and guanine
CC: corpus callosum
CV: coefficient of variation
DBS: disease burden score
DCL: diagnostic confidence level
DT-MRI: diffusion tensor magnetic resonance imaging
FDM: frequency difference mapping
HD: Huntington’s disease
HTT: huntingtin
mGRE: multi-echo gradient-recalled echo
MoCA: Montreal Cognitive Assessment
MTR: magnetization transfer ratio
MW: myelin water
MWF: myelin water fraction
MWI: myelin water imaging
PCA: principal component analysis
PEBL: Psychology Experiment Building Language (PEBL)
RF: radiofrequency
TE: echo time
TMS: total motor score
TOPFUK: Test of Premorbid Functioning - UK Version
TR: repetition time
UHDRS: Unified Huntington’s Disease Rating Scale
WAIS-R: Wechsler Adult Intelligence Scale-Revised
WM: white matter

## Funding & Acknowledgements

The present research was funded by a Wellcome Trust PhD studentship to CC (ref: 204005/Z/16/Z); DKJ and EK were supported by a New Investigator Award (to DKJ) from the Wellcome Trust (ref: 096646/Z/11/Z) and a Strategic Award from the Wellcome Trust (ref: 104943/Z/14/Z). We thank Dr Slawomir Kusmia and Dr Mark Drakesmith for their support with the project.

## Appendix A: Details on mGRE signal pre-processing

In this work the mGRE signal was acquired using bipolar gradient readouts with the polarity of the first echo inverted halfway through the number of repeats. We use ± sign to indicate where the polarity of the read gradient may affect the corresponding mGRE signal. The complex mGRE signal was reconstructed per receive channel *k* and gradient echo *n* at corresponding echo time TE_*n*_ and can be represented as follows (Sati et al., 2013; Tendler & Bowtell, 2019; Thapaliya et al., 2017; Wharton & Bowtell, 2012; Nam et al. 2015a,2015b; Kleban et al. 2021):

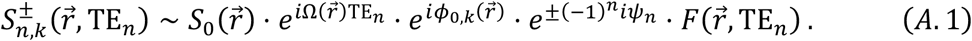

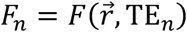 represents the complex WM signal as a function of echo time, which in this work was modelled using a three-pool model.

The exponentials preceding *F_n_* in Equation (A.1) reflect macroscopic spatio-temporal phase effects which contribute to the total complex mGRE signal and need to be addressed before *F_n_* can be analysed (Lee et al., 2018; Sati et al., 2013; Tendler & Bowtell, 2019; Thapaliya et al., 2017; Wharton & Bowtell, 2013):

1. The frequency Ω represents the effect of large-length-scale field perturbations; 2) The phase *ϕ*_0,*k*_ describes collective phase offsets including those specific for each receiver channel; 3) The phase *ψ_n_* highlights phase differences arising when signal from the same echo is acquired with opposite gradient polarities.

We correct for the phase effects in following steps:

1. The phase shifts *ψ_n_* can be removed by multiplying 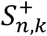 by 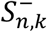 (Kleban et al. 2021):

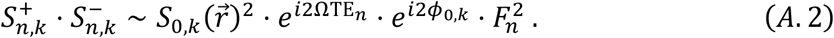
2. Remove phase offsets, particularly those varying between channels (Tendler & Bowtell, 2019; Kleban et al. 2021; Bydder et al. 2002):

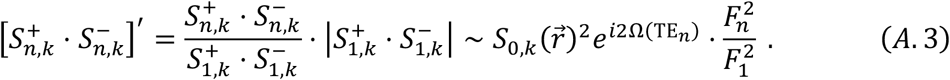

This is the first step of frequency difference mapping which in this case also removes phase inconsistencies between the receive channels, so that the coil combination can be applied next.
3. The signals from all receive channels are combined by calculating the sum of over the channels weighted by the channels’ sensitivity *m_k_* (Bydder et al., 2002; Roemer et al., 1990).

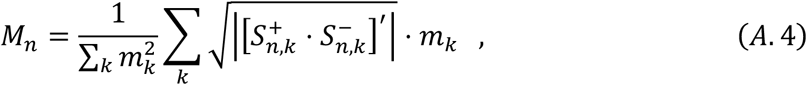

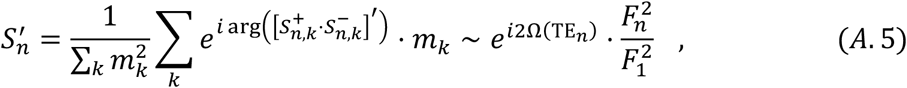

where *m_k_* was approximated from signal magnitude of the first echo:

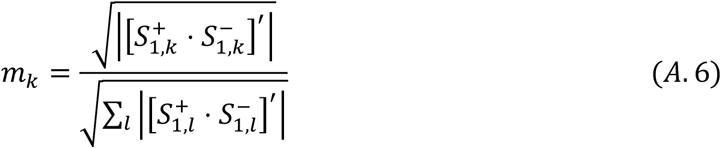
4. The final step of FDM calculation aims to remove the effects of Ω (Tendler & Bowtell, 2019; Kleban et al. 2021):

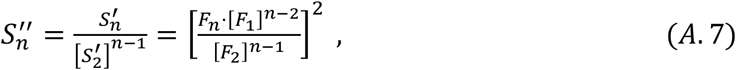

which can be translated to frequency:

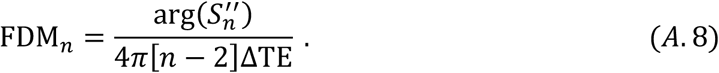

As evident from equation A.7, each echo contributes to the estimation of the frequency difference.

## Appendix B: Simulations to address the impact of 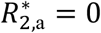 assumption on MW signal fraction

We simulated the complex mGRE signal using a 3-pool model:

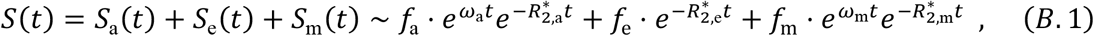

and the signal parameters reported by Sati et al (2013), also listed in Table B1, unless stated otherwise. We then simulated two cases and varied: i. the myelin water signal fraction, *f*_m_, from 0.1 to 0.2; or ii. the intra-axonal relaxation rate, 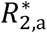, from 15 to 35 s^-1^. Figures 1(2)A and B show the frequency difference and the magnitude of the signal for each case, respectively. We then fitted the 3-pool model assuming 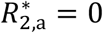, to test the influence of this assumption on the estimated myelin water signal fraction. Figure 1C shows the estimated MW signal fraction plotted against the input value. We also tested the effect of the assumption 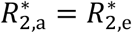 in Figure 1D. Finally, Figure 2C shows MW signal fraction values estimated under the assumption of 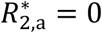 and plotted against the input 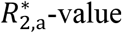. Colormaps in Figures B1 and B2 reflect the variable input values for *f*_m_ and 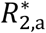, respectively.

Overall, though a small bias can be observed in the estimated MW signal fractions for the given extra-axonal R2*, such bias is of the same magnitude for all input MW signal fractions and is smaller than the calculated CV values for this segment of the CC, suggesting that any bias introduced by possible differences in axonal R2* between groups is unlikely to have affected results in our study.

**Table B1.** Input parameters for the simulated signal. The MW signal fraction f_m_ was varied from 0.1 to 0.2 for Figure B1, and the intra-axonal relaxation rate was varied from 15 to 35 s^-1^ for Figure B2.

| $\omega_a$ [Hz] | $\omega_m$ [Hz] | $R_{2,m}^*$ [$s^{-1}$] | $R_{2,a}^*$ [$s^{-1}$] | $R_{2,e}^*$ [$s^{-1}$] | $f_a/f_e$ | $f_m$ |
| --- | --- | --- | --- | --- | --- | --- |
| -6 | 30 | 160 | 25 | 40 | 2/3 | 0.15 |

**Figure B1.**
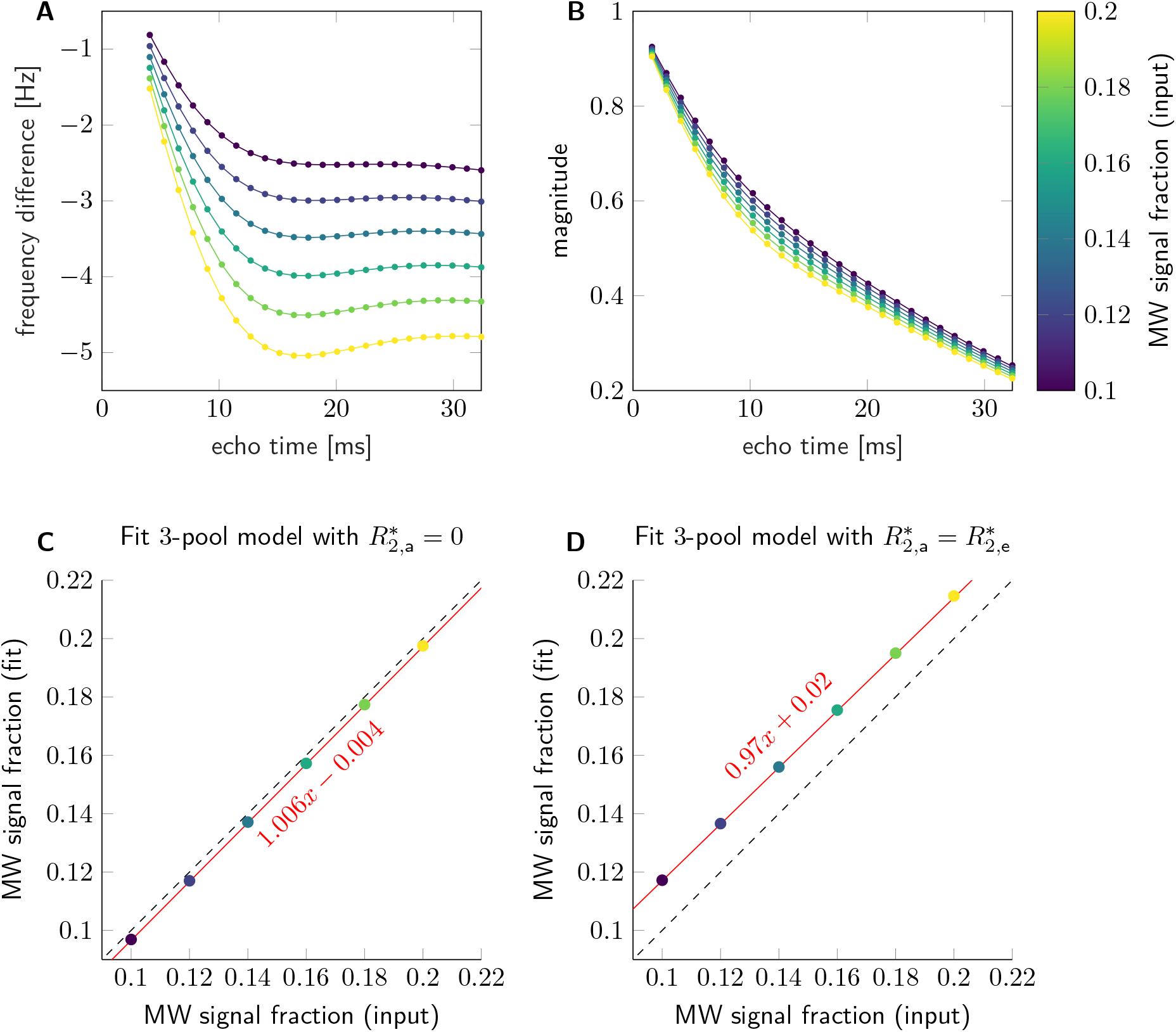
Frequency difference (A) and magnitude (B) were estimated from a complex signal, which was simulated for myelin water signal fraction, f_m_, ranging from 0.1 to 0.2. Other parameters used to simulate the 3-pool complex signal are listed in Table 1. The colour bar corresponds to f_m_. The 3-pool model was then fitted for two assumptions: 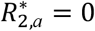 and 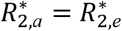, and the estimated MW signal fraction values were plotted against the input values in (C) and (D), respectively. The red line represents the linear relationship between the estimated f_m_ and the input f_m_, slopes and intercepts are indicated in the plots.

**Figure B2.**
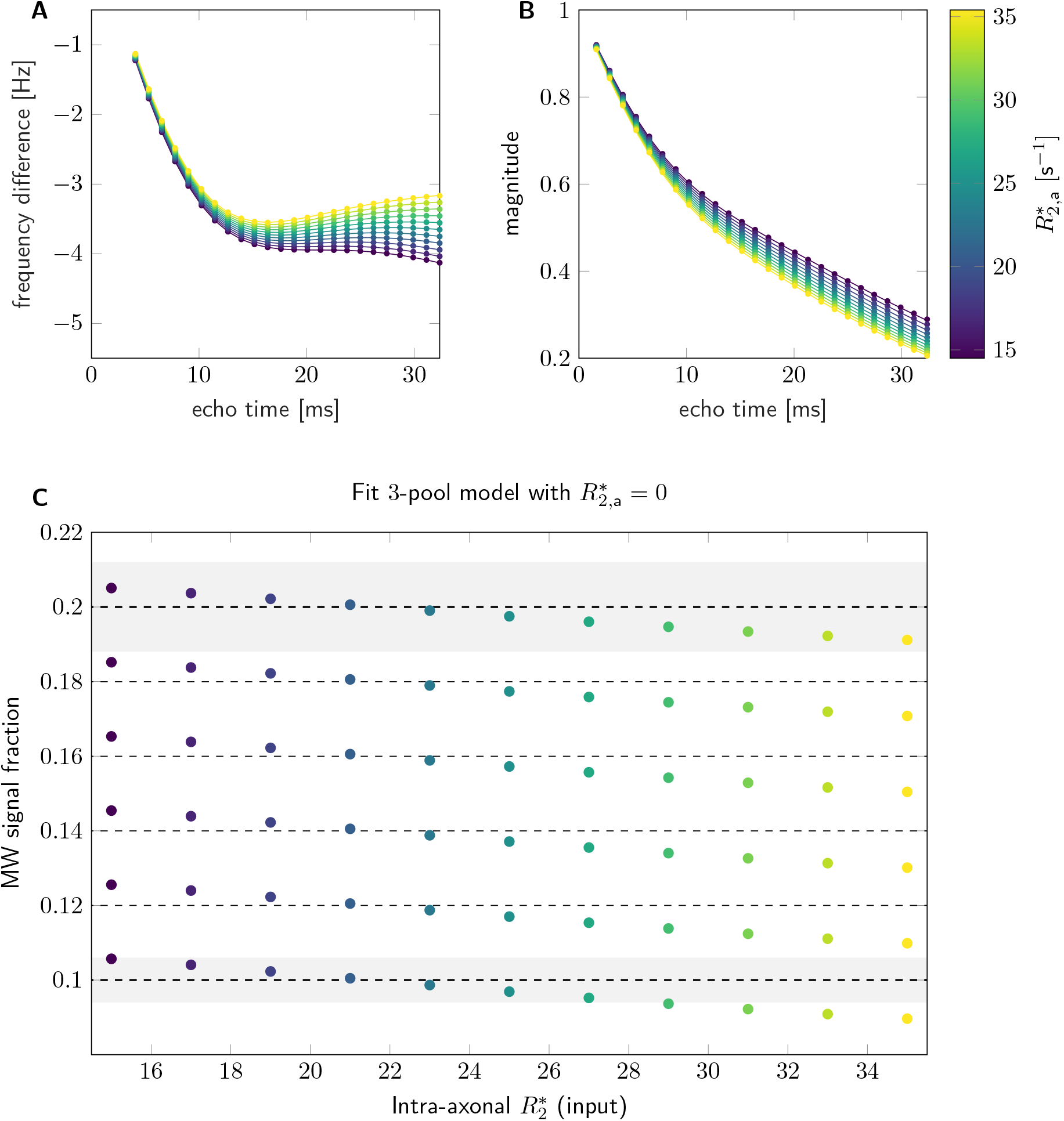
Frequency difference (A) and magnitude (B) were estimated from a complex signal, which was simulated for intra-axonal relaxation rate 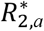 ranging from 15 to 35 s^-1^. Other parameters used to simulate the 3-pool complex signal are listed in Table 1. In C, MW signal fraction was estimated under assumption 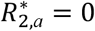 and plotted against the input intra-axonal relaxation rate. Dashed lines indicate the input MW signal fractions. Gray areas are equivalent to 12% (±6%) of 0.1 and 0.2, respectively. The value of 12% is equivalent to coefficient of variation of f_m_ in the splenium of the corpus callosum estimated in the reproducibility study (cf., Table 6). The colour bar corresponds to 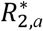.

